# Compass orientation of a migratory mammal in total darkness

**DOI:** 10.64898/2026.10.07.757263

**Authors:** Fyodor Cellarius, Jonas Beer, Konrad Sucher, Agate Ozoliņa, Viesturs Vintulis, Andreas Jechow, Oliver Lindecke

**Affiliations:** Neurosensorics/Animal Navigation Group, Institute for Biology and Environmental Sciences, Carl-von-Ossietzky Universität Oldenburg, Carl-von-Ossietzky-Straße 9-11, 26129 Oldenburg, Germany; Quantum and Environmental Photonics Group, Brandenburg University of Applied Sciences, Magde-burger Str. 50, 14770 Brandenburg an der Havel, Germany; Bat Research Society of Latvia, "Niedrāji", Glūdas pagasts, Jelgavas novads, LV–3040, Latvia; Laboratory of Ornithology, Institute of Biology, University of Latvia, Jelgavas Street 1, Rīga LV–1004, Latvia

**Keywords:** bat migration, cryptochrome, magnetoreception, *Pipistrellus nathusii*, radical pair mechanism

## Abstract

Many animals use the Earth’s magnetic field to orient and navigate over long distances, yet the sensory mechanism that underlies this ability remains unresolved. A central question is whether the compass depends on light: a cryptochrome-based radical-pair mechanism, best characterised in birds, requires a photon to generate the magnetically sensitive species, whereas magnetic particle-based and inductive mechanisms do not. In mammals, which lack the light-sensitive cryptochromes that the avian model requires, this question is unresolved. Here we tested whether migratory bats, *Pipistrellus nathusii*, require light for migratory orientation, studying wild migrants at a major flyway under laboratory-controlled light conditions, including verified total darkness. Bats were acclimated to starlight-level illumination or to total darkness before release into a circular arena in which take-off direction was recorded, and take-off latency was taken in every trial as a control for behavioural state. Bats released under starlight-level illumination, and bats released in total darkness after starlight acclimation, both oriented in the south-westerly direction typical of free-flying bats at this flyway; their mean headings lay 11° apart and did not differ in direction, in angular concentration, or in take-off latency. Bats released in total darkness after prolonged darkness produced the same mean heading but did not reach significance under a Rayleigh test, and did not differ from either other group. To our knowledge, this is the first demonstration that a migratory mammal expresses its seasonally appropriate migratory heading in total darkness. Bats orienting in darkness took off five to eight minutes after their last photon exposure, seven to eight orders of magnitude beyond the microsecond lifetime of the only cryptochrome radical pair demonstrated to exist and to be magnetically sensitive. Because a compass reading is informative only at the moment of the directional decision, a signal generated before a bat entered darkness could not encode its heading at take-off; the magnetically sensitive species must therefore form without photons. The radical pairs capable of doing so have not been shown to exist, even *in vitro*. A migratory mammal thus expresses its seasonal heading without concurrent light, in a lineage lacking the photoreceptor on which the light-dependent compass in birds relies.

## Background

Magnetoreception is the only vertebrate sensory modality for which no primary receptor has been identified [1]. Three mechanisms remain viable hypotheses, and they separate along a single axis: whether the primary detection step requires light. In the radical-pair mechanism, absorption of a photon by a flavin-binding cryptochrome generates a pair of radicals whose spin dynamics, and therefore whose reaction yield, depend on the orientation of the ambient magnetic field [2]; it is supported *ex vivo* by the magnetic sensitivity of cryptochrome 4 from a night-migratory songbird [3]. In the magnetic-particle mechanism, ferrimagnetic crystals transduce the field mechanically, as they do in magnetotactic bacteria [4,5]; the hypothesis has been proposed for mammals [6,7] and has behavioural support in a bat, whose homing orientation is altered by a magnetic pulse [8]. In the induction mechanism, movement of the head through the geomagnetic field drives currents in the fluid-filled semicircular canals, read out by hair cells bearing electroreceptive voltage-gated ion channels [9]; a whole-brain screen in pigeons found a vestibular-mesopallial circuit activated by magnetic stimuli equivalently in light and in darkness [10]. Neither of the latter two mechanisms requires photons. Whether magnetic orientation can be expressed in darkness has therefore been treated as the sharpest behavioural discriminant available between the light-dependent candidate and the light-independent ones.

Bats are capable of performing long-range migration, with the Nathusius’ pipistrelle (*Pipistrellus nathusii*) holding records of more than 2,000 km [11,12]. However, the mechanisms by which migratory mammals navigate over such distances remain unknown. The set of cues available for long-range navigation over thousands of kilometres is thought to be limited to large-scale landmarks (e.g., coastlines, mountain ranges) and the Earth’s magnetic field. The latter is the best candidate to explain bats’ ability to maintain a compass direction during migration [13], as they can consistently demonstrate migratory orientation even under enclosed, controlled conditions [14]. Indeed, bats are known to use the magnetic field as a source of directional information [15]. Earlier work on the Chinese noctule (*Nyctalus plancyi*) further reported polarity-based roosting preferences in small groups of captive bats held in total darkness [16], with the authors arguing from cave-roosting ecology that a light-based receptor was unlikely. That study tested resting alignment rather than a migratory compass, did not manipulate light as an experimental variable, and relied on extended exposures of the same small cohort across conditions; they therefore leave open whether the migratory compass orientation itself requires light. However, it has been demonstrated that pipistrelles (*Pipistrellus pygmaeus*) integrate magnetic inclination (the angle between the magnetic field vector and the Earth’s surface) to orient during migration [17].

Darkness is nevertheless a blunter instrument than it first appears. A light-initiated signal need not be consumed at the instant it is created: during the re-oxidation of the fully reduced flavin, a second radical pair (RP2) can form without photon input, on a timescale of minutes rather than the microseconds of primary radical-pair recombination, so magnetic responses observed in darkness cannot in themselves be read as evidence against radical-pair involvement [1]. Longer-lived variants have been reported: in the firebug *Pyrrhocoris apterus*, which expresses only type II cryptochrome, a cryptochrome-dependent magnetic response conditioned under light survived at least 24 h of darkness [18], although the magnetic sensitivity of superoxide-containing radical pairs is contested on physical-chemistry grounds [19]. What a darkness experiment can establish is therefore not light-independence, but whether the magnetically sensitive species must be generated photochemically at the moment the field is read. We designed this study around that distinction.

The case for a light-dependent compass rests almost entirely on birds, and the molecular basis established there cannot simply be assumed to transfer. Mammals express only type II cryptochromes (CRY1 and CRY2), which lack the canonical photoreceptive function required to generate a light-driven radical pair [20], so whether a light-dependent compass is available to mammals at all remains open. A heterologous assay reported that human CRY2 restores light-dependent magnetosensitivity in cryptochrome-null Drosophila [21], indicating that a mammalian type II cryptochrome can in principle support such a response in an invertebrate background. This does not establish a native function in mammals, and the Drosophila behavioural paradigm on which the claim rests has not withstood high-powered replication [1,22]. Magnetic orientation is nevertheless well documented in mammals, including bats: big brown bats *(Eptesicus fuscus*) are disoriented by a rotated field [15], and migratory soprano pipistrelles (*Pipistrellus pygmaeus*) resolve magnetic inclination during a defined compass-calibration window [17]. Which mechanism underlies the mammalian compass is unknown, and because mammals lack the molecule on which the avian model depends, they are the lineage in which the light question is both most open and most diagnostic.

In birds, the principal behavioural signature of radical-pair involvement is the disruption of magnetic compass orientation by radiofrequency (RF) electromagnetic fields [23–25]; RF exposure has recently been reported to disrupt migratory orientation in *P. pygmaeus* as well [14]. Whether that signature is diagnostic in mammals is less clear, because CRY4, the retinal cryptochrome that carries the avian radical pair, is absent in mammals [20]. RF sensitivity in a mammal is therefore consistent with radical-pair involvement without establishing it. The caution is not peculiar to RF: an animal may carry more than one magnetosensor [26–29], so disrupting one can disorient without implicating a particular mechanism, and non-specific effects are hard to exclude [30]. None of this, however, bears on whether the mechanism requires light. The hybrid mechanism proposed by Hore [31], in which a single-domain magnetite particle amplifies the field experienced by a nearby radical pair, sharpens the point: a sensor containing magnetite may still require light. Testing the light requirement probes a mechanistic necessity rather than a disruption signature.

Migratory bats offer a tractable mammalian model. Nathusius’ pipistrelle (*P. nathusii*) migrates over 2,000 km between the Baltic and south-western Europe [11,12], and its seasonally appropriate autumn departure direction at the Baltic flyway is established independently of any arena, by radio-tracking, by releases after translocation [13,32], and by ring recoveries showing that bats ringed at this site are recovered to the south-west [33]. Take-off direction from a circular release box reproduces migratory heading of pipistrelles under enclosed conditions [14,34], and migratory *P. pygmaeus* tested with the assay at the same site, shifted take-off direction when magnetic inclination is manipulated during the compass-calibration window [17]. We therefore treat the heading recorded in this assay as the output of a magnetically informed compass, an assumption we examine explicitly in the Discussion. Because the assay reads a direction rather than a position, it addresses compass orientation and not map-based navigation.

We tested wild migratory Nathusius’ pipistrelle bats, *Pipistrellus nathusii*, under three conditions: star-light-level illumination before and during the test (control); total darkness at test following starlight acclimation; and total darkness at test following prolonged total darkness. Light was calibrated to the irradiance of a natural moonless night sky, and the darkness conditions were verified photometrically (**Fig. 1**). A magnetic compass that operates in darkness is, in itself, compatible with a magnetite-based mechanism. For reference, the avian light-dependent compass has been behaviourally demonstrated at light levels down to those of civil twilight [35]; the conditions imposed here, the light of a moonless starry night sky and verified total darkness, lie below that range. We do not assume the mammalian mechanism to be homologous with the avian one, and additional measurements verified that darkness treatments fell below the visual detection threshold of the experimental bats (“total darkness”). The design yields two single-factor contrasts. Comparing the starlight control with bats released in darkness after starlight acclimation, with the preceding light history held constant, tests whether the compass requires photons at the moment the field is read. Comparing the two darkness groups, with the test condition held constant, varies the interval since the last photon exposure by more than an order of magnitude. Because light also modulates emergence and flight activity in this species independently of magnetoreception [36–38], we recorded take-off latency in every trial as a control for behavioural state

**Figure 1.**
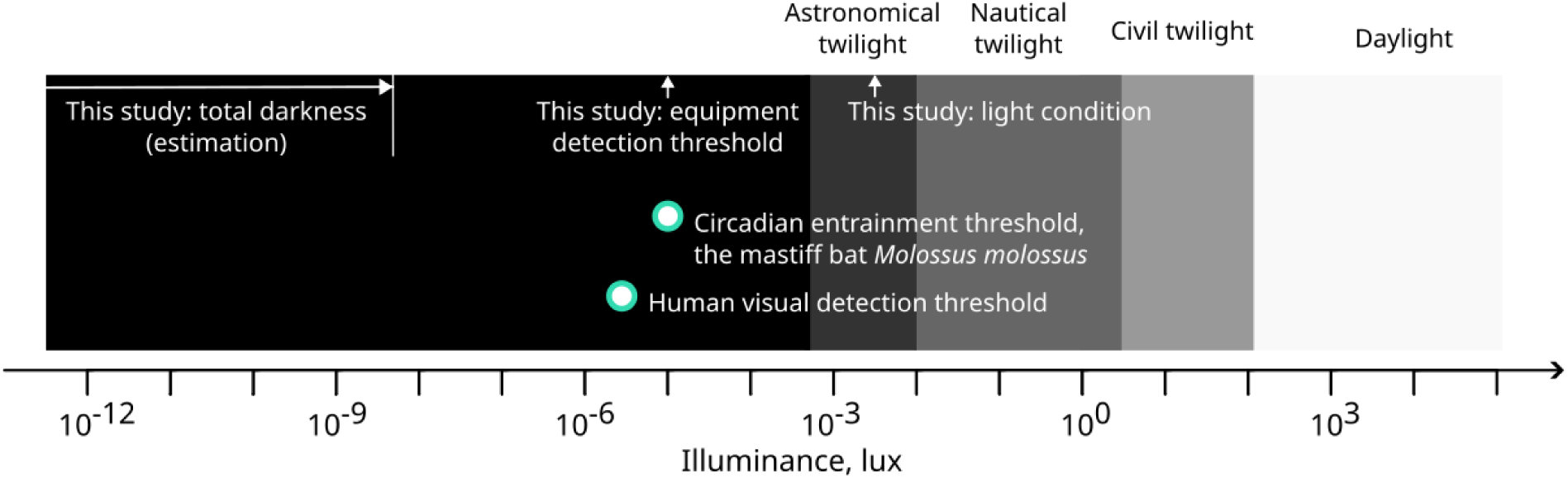
Experimental light conditions in the context of global illuminance thresholds. Light conditions in this study corresponded approximately to a starry moonless night / astronomical twilight [39]. The total darkness condition inside the yurt was below 70 plx and the upper short-term exposure below 5 nlx; it lies below the human visual detection threshold [40] and below the lowest known circadian photic-entrainment threshold of a mammal, recorded for the mastiff bat *Molossus molossus* [41].

## Results

Bats in all experimental conditions exhibited south-westerly mean orientations. The control group, exposed to light matching the natural night-sky both before and during testing, showed a significant unimodal south-westerly orientation (α = 223°, N = 23, r = 0.44, Rayleigh p = 0.01; **Fig. 2A**). A similar unimodal pattern was observed in bats tested in total darkness following starlight acclimation in the pre-exposure period (α = 234°, N = 24, r = 0.40, Rayleigh p = 0.02; **Fig. 2B**). Bats acclimated to total darkness and released in total darkness exhibited a similar mean direction (α = 235°, N = 25; r = 0.24; **Fig. 2C)**, although their orientation did not reach statistical significance (Rayleigh p = 0.23). A maximum likelihood analysis indicates that the alternatives supported for this sample include both uniform and unimodal (‘oriented’) models (models with ΔAICc < 2; **Table S1,** Electronic supplementary material 1). An estimation of Type II error rate for Rayleigh test shows the rate of approximately 0.69 for the samples drawn from a Von Mises distribution with a concentration parameter analogous to the dark acclimated sample (**Fig. S1,** Electronic supplementary material 1). Pairwise comparisons revealed no detectable differences between any pair of groups in mean direction or in angular concentration. Neither darkness group differ from the control (starlight acclimation: MWW p = 0.99, concentration test p = 0.81; dark acclimation: MWW p = 0.70, concentration test p = 0.30), and the two darkness groups did not differ from one another (MWW p = 0.76, concentration test p = 0.42; **Fig. 2**). A post hoc sensitivity analysis showed that, if the dark-acclimated sample were uniformly distributed, the probability of obtaining a MWW test p-value equal to or greater than the observed was low (control versus dark acclimation: 0.02; light versus dark acclimation: 0.01), indicating that the lack of significant differences likely reflects similarity between the samples rather than low statistical power. Take-off latency did not differ across conditions (Kruskal-Wallis H = 1.76, p = 0.41; control 104 ± 48 s, starlight acclimation 90 ± 45 s, dark acclimation 85 ± 55 s); bats acclimated to darkness took off, if anything, sooner than controls, giving no indication that darkness slowed the initiation of flight. Nor did the proportion of bats failing to take off within the 3 min window differ across conditions (control 3 of 26; starlight acclimation and take-off in total darkness 0 of 24; total dark acclimation and take-off 3 of 30; Monte Carlo Fisher’s exact test p = 0.28; control versus dark acclimation, Fisher’s exact test p = 1.00; 2 animals from the dark-acclimated group are included here as they successfully took off, but excluded from the orientation analysis since they did not leave a distinguishable track for orientation measurement), so the analysed samples were not differentially selected. The duration of acclimation was not fixed: bats were placed in the acclimation boxes in a single session and tested sequentially, so the interval before take-off ranged from approximately 90 min to approximately 6.5 h; take-off direction was independent of this interval in every condition, expressed as hours after sunset (circular-linear correlation: control r = 0.31, p = 0.33; starlight acclimation r = 0.23, p = 0.53; dark acclimation r = 0.20, p = 0.60), so orientation did not decay over the range of intervals imposed.

**Figure 2.**
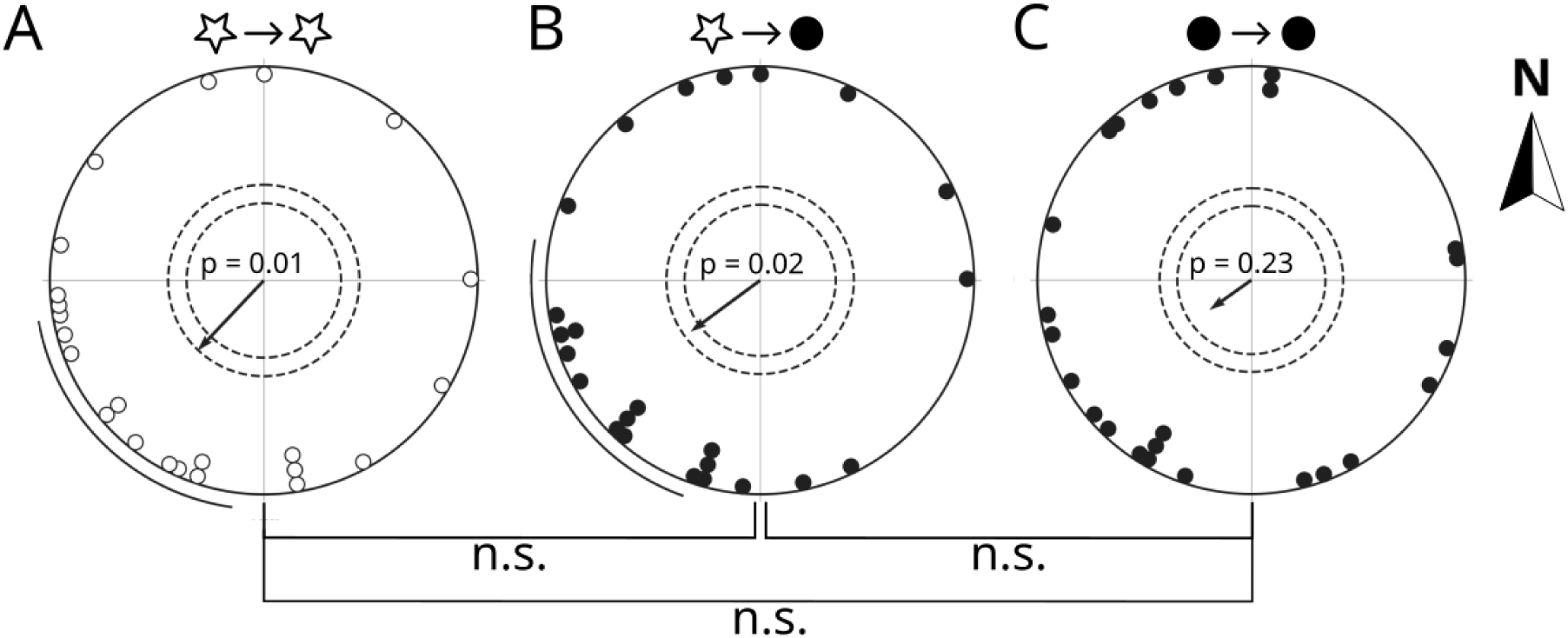
Orientation of migratory *P. nathusii* released in the local geomagnetic field under starlight and total darkness conditions. A) Control bats, exposed to light both before and during testing, showed significant south-westerly orientation (α = 223°, N = 23, r = 0.44, Rayleigh p = 0.01). B) Bats tested in total darkness following starlight acclimation performed similarly to controls (α = 234°, N = 24, r = 0.40, Rayleigh p = 0.02); the two groups had identical pre-exposure light conditions up to release and differed only in whether photons were available during the take-off decision. C) Bats tested in total darkness after prolonged dark acclimation produced the same mean heading but did not reach significance (α = 235°, N = 25; r = 0.24; Rayleigh p = 0.23). All bats were transferred to the acclimation boxes under indirect daylight, identical across conditions, before the acclimation in pre-exposure under light or total darkness began; the acclimation interval ranged from approximately 90 min to 6.5 h, and a 2 min settling period in an opaque box preceded release in all conditions. Each point represents the take-off orientation of one bat. Outer segments on A) and B) correspond to 95% confidence intervals for the mean direction. The arrow indicates the mean vector, and the inner dotted circles correspond to 0.01 and 0.05 significance levels of the Rayleigh test.

## Discussion

Migratory *Pipistrellus nathusii* bats oriented in a seasonally appropriate compass direction in total darkness under laboratory conditions. Removing light at the time of testing did not alter compass orientation. Bats released in total darkness took off within 11° of the mean heading of controls under starlight-level illumination, with statistically indistinguishable mean direction and angular concentration, suggesting that the same compass system was read out under both conditions rather than a distinct response engaged only in darkness. In birds tested in total darkness, by contrast, an orientation response is reported that does not correspond to the seasonally appropriate migratory direction; that response seems to be under magnetic control but does not engage the light-dependent inclination compass, and it was attributed to a separate receptor and termed a "fixed-direction" response [42]. Bats both tested and acclimated to total darkness before release produced a mean heading in the same south-westerly direction (**Fig. 2C**), but did not reach significance, and the maximum-likelihood analysis gives equal support to random and oriented alternatives. However, a complete loss of direction appears unlikely based on the detailed comparison of the two darkness groups, and we therefore do not treat that group as evidence of a light requirement. To our knowledge, no mammal has previously been shown to express a seasonally appropriate migratory heading in total darkness: magnetic responses in darkness are documented in mole-rats [43] and as polarity-dependent roosting alignment in a bat [16], but neither is a migratory heading, and among vertebrates a migratory heading in darkness is otherwise known only from hatchling sea turtles [44,45] and sockeye salmon [46], while laboratory experiments in zebrafish and medaka provide independent evidence that teleosts can respond directionally to magnetic fields without visible light [47].

The persistence of compass orientation under total darkness is difficult to reconcile with a magnetoreceptor that requires light when it senses the magnetic field, i.e., photon input at the moment of transduction. The canonical radical-pair mechanism, as characterised in bird research, is initiated when a cryptochrome flavin (FADox) absorbs a photon, generating a light-driven radical pair (RP1) whose spin-state dynamics are modulated by the ambient magnetic field and transduced into a directional signal [2,3]. For this pathway, light must be available at the moment of transduction. The interval involved is extreme: the only cryptochrome-derived radical pair demonstrated to exist and to generate magnetic field effects, [FAD•− TrpH•+], is formed photochemically and has a lifetime of the order of one to five microseconds [48], yet bats released in total darkness took off five to eight minutes after their last photon exposure, seven to eight orders of magnitude later. A radical pair generated before the animal entered darkness cannot account for this, and not only because it would have decayed: a compass reading encodes the direction of the field relative to the head at the moment it is taken, so a signal generated minutes earlier, while the bat was handled, carried and confined, encodes a heading it no longer holds. To use it, the bat would have to have taken a bearing at the last moment of illumination and maintained it by path integration through three minutes in a cloth bag, two minutes in an opaque box and its own movements in the arena, which we regard as implausible. The magnetically sensitive species must therefore be generated in darkness. The present results are therefore inconsistent with a compass requiring photon input at transduction (the canonical RP1 pathway) in *P. nathusii*; we return below to the specific variant of the hypothesis in which a photo-initiated signal persists into darkness. In birds, orientation can persist in total darkness but has been reported as a polar "fixed-direction" response that does not follow the migratory direction and does not engage the light-dependent magnetic inclination compass. Hence, Stapput et al. (2008) hypothesize that a second, light-independent receptor plays a role in birds [42]. However, the seasonally appropriate south-westerly heading maintained by our bats, stable across all three test conditions, is therefore unlike this avian darkness response. The south-westerly headings recorded here coincide with the seasonally appropriate departure directions of free-flying *P. nathusii* at this flyway, both radio-tracked during autumn migration and released after translocation [13,32], indicating that the arena captures the species’ genuine migratory orientation rather than an arbitrary heading. The rotational symmetries of the test room and the arena, and the randomised entry direction argue against a fixed directional cue within the apparatus.

A photo-initiated signal could, in principle, sustain cryptochrome-dependent magnetoreception transiently in darkness through the reoxidation radical pair (RP2), formed between the reduced flavin and superoxide during a re-oxidation step that requires no photons and proceeds over minutes [1]. This is the only remaining radical-pair account of our result, and it is a demanding one. Of the two candidate reoxidation pairs, [FAD•− Z•] and [FAD•− Asc•−], it has been noted that their existence, let alone their magnetic sensitivity, has not been demonstrated even in vitro [48], and the viability of superoxide-containing radical pairs as magnetoreceptors has been questioned on on physical-chemistry grounds [19,31]. The hybrid mechanism, in which a magnetic particle amplifies the field experienced by a nearby radical pair [31], does not resolve this: the amplification increases sensitivity, but the radical pair must still be generated photochemically. The interval that our data require this signal to bridge is not, in any case, of the order of minutes: bats acclimated to darkness took off approximately 1.5–6.5 h after their last exposure to light, without their take-off orientation becoming uniform. One report in an invertebrate with a mammalian-type cryptochrome is worth noting: in the firebug *Pyrrhocoris apterus*, which expresses only Cry II, a Cry II-dependent magnetic response conditioned under light has been reported to persist for at least 24 h in total darkness and to be lost in a Cry II null mutant [18]. Our dark-pretreated group is not a clean test of this scenario. We therefore constrain rather than exclude these persistence-based variants: mechanisms requiring signal half-lives on the order of minutes are difficult to reconcile with any of our groups, whereas mechanisms with half-lives well beyond 6.5 h remain formally possible.

Three candidate mechanisms remain consistent with our result: a radical-pair mechanism, a magnetic-particle mechanism, and electromagnetic induction. Our behavioural data do not rank them, and the mag-neticparticle and radical-pair hypotheses are not mutually exclusive: in the hybrid mechanism proposed by Hore [31], a single-domain magnetite particle amplifies the geomagnetic field for a nearby radical pair, so a sensor may contain magnetite yet remain light-dependent. The magnetic-particle hypothesis has behavioural support in a bat, where magnetic pulses altered orientation in *Eptesicus fuscus* [8]; pulse remagnetisation is a mechanistically distinct diagnostic that could be applied to *P. nathusii*. The electromagnetic induction hypothesis has recently gained empirical traction: Nimpf et al. [9] proposed that pigeons detect magnetic fields via induction in the fluid-filled semicircular canals of the inner ear, transduced by hair cells expressing the long isoform of the voltage-gated calcium channel Ca_V1.3, the same channel that mediates electroreception in elasmobranchs. A subsequent whole-brain neuron activity screening in pigeons found that a vestibular-mesopallial circuit is activated by magnetic stimuli equivalently in light and darkness [10], consistent with, but not yet proof of, a light-independent inner-ear magnetosensor. This model has, however, recently been challenged on physical grounds: an information-theoretic analysis concludes that Faraday induction in semicircular canals of pigeon dimensions does not carry sufficient directional information to support a compass [49], a constraint that would apply more strongly to bats such as *P. nathusii*, whose canals are smaller. This reduces, but does not eliminate, the plausibility of an inductive mechanism. None of these mechanisms has been demonstrated in a mammal, and no mammalian magnetoreceptor has been identified. These results constrain the receptor: it must yield a compass bearing without concurrent light, in a lineage that lacks the photoreceptive cryptochrome on which the avian model rests. Orientation over long distances in darkness is an ordinary circumstance for a range of mammals beyond bats, among them whales and seals, which spend hours at a time in near-darkness at depth in the ocean basins [50–52]. Bats offer a tractable model to understanding how a compass that holds a heading into the dark, once light has been available, could have evolved wherever that transition is routine. Magnetic orientation in mammals has so far been examined in a handful of species and traced to a receptor in none of them; in a migratory bat it can now be studied in a wild animal performing, on the night it is tested, the task for which its compass is used.

## Materials & Methods

### Study site and experimental animals

The experiments were conducted in the period from 21.08 to 01.09.2025 at Pape Ornithological Station, Latvia. In total, 81 adult migratory pipistrelles were captured with a large funnel trap. The bats’ overall fitness, sex and age were determined on the capture night. Between capture and testing, bats were held in groups of up to 5 individuals in a windowless wooden hut in wooden boxes with a sliding door and ventilation apertures; the hut was dark except when entered by an experimenter. On the night of testing, bats were handled individually by the same experimenter and transferred to the acclimation boxes between 21:00 and 21:30, in a room lit only by indirect daylight through a single-glazed, east-facing window, approximately 2 m from it and not in direct view of it; no artificial lighting or head torches were used. This was the last exposure to broad-spectrum light before take-off, common to all three exposure conditions. Bats kept for longer than one day after capture were fed and offered water in the evenings every day until release.

### Experimental design

Three groups of bats were tested for take-off orientation: controls, acclimated and tested under starlight illumination; light-acclimated bats tested in total darkness; and dark-acclimated bats tested in total darkness. All treatments consisted of the same functional blocks (**Fig. 3**).

**Figure 3.**
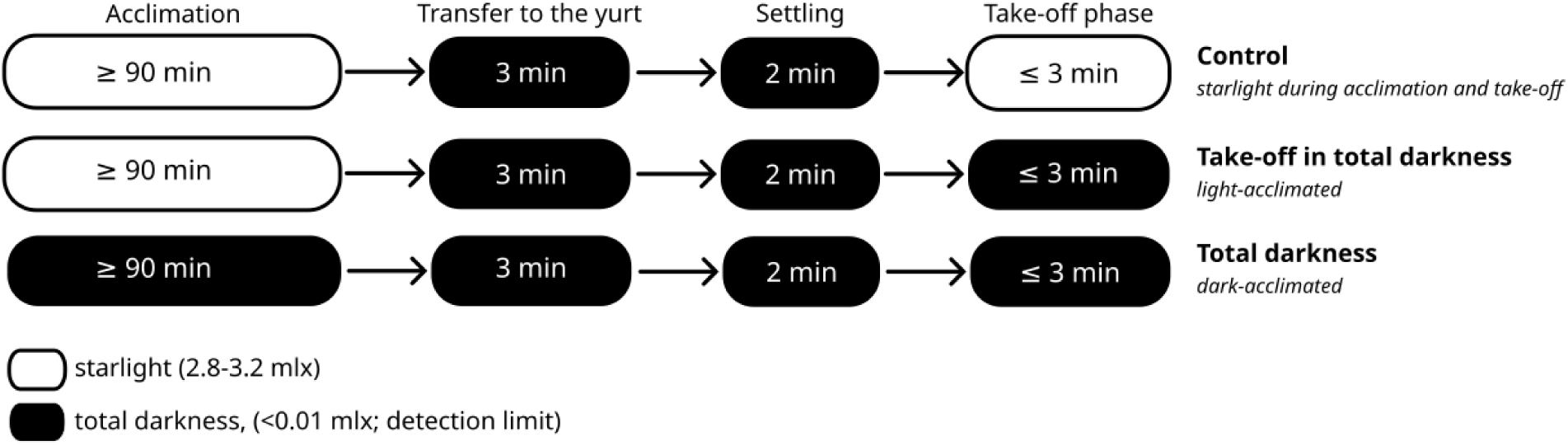
Experimental design. Bats were acclimated under either starlight illumination or total darkness. After acclimation, all three groups underwent the same transfer and settling procedure in total darkness. Take-off orientation was then recorded under either starlight illumination or total darkness.

Initially, bats were acclimated either under starlight illumination or in total darkness. Because all bats were placed in the acclimation boxes during a single session and tested sequentially, acclimation duration ranged from a minimum of 90 min for the first animal tested to 387 min for the last, corresponding to approximately 1.5–6.5 h of pre-exposure. Take-off direction was independent of this interval within each condition (see Results), so acclimation duration was not treated as an experimental variable.

After acclimation, bats were transferred from the acclimation chamber (see Experimental procedure) to the testing area in total darkness. The transfer took 3 min and was followed by a 2-min settling period, during which the bat remained inside the orientation arena under a settling box positioned in the centre of the circular arena. All bats, including controls, therefore spent 2 min in total darkness immediately before release. Because this interval was identical across treatments, the control and light-acclimated total-darkness groups differed specifically in the presence or absence of light during the subsequent take-off phase. After settling, each bat was given up to 3 min to take off.

### Experimental procedures

Bats were tested between 22:00 and 04:00 inside a Mongolian yurt, 5.6 m in diameter, with a height of 2.4 m at the centre and 1.5 m at the edge (**Fig. 4A**), with a Circular Release Box – a circular arena used to assess a bat’s orientation in an enclosed space – placed at the centre. The interior of the yurt is rotationally symmetric, which prevents animals from obtaining directional information via echolocation.

**Figure 4.**
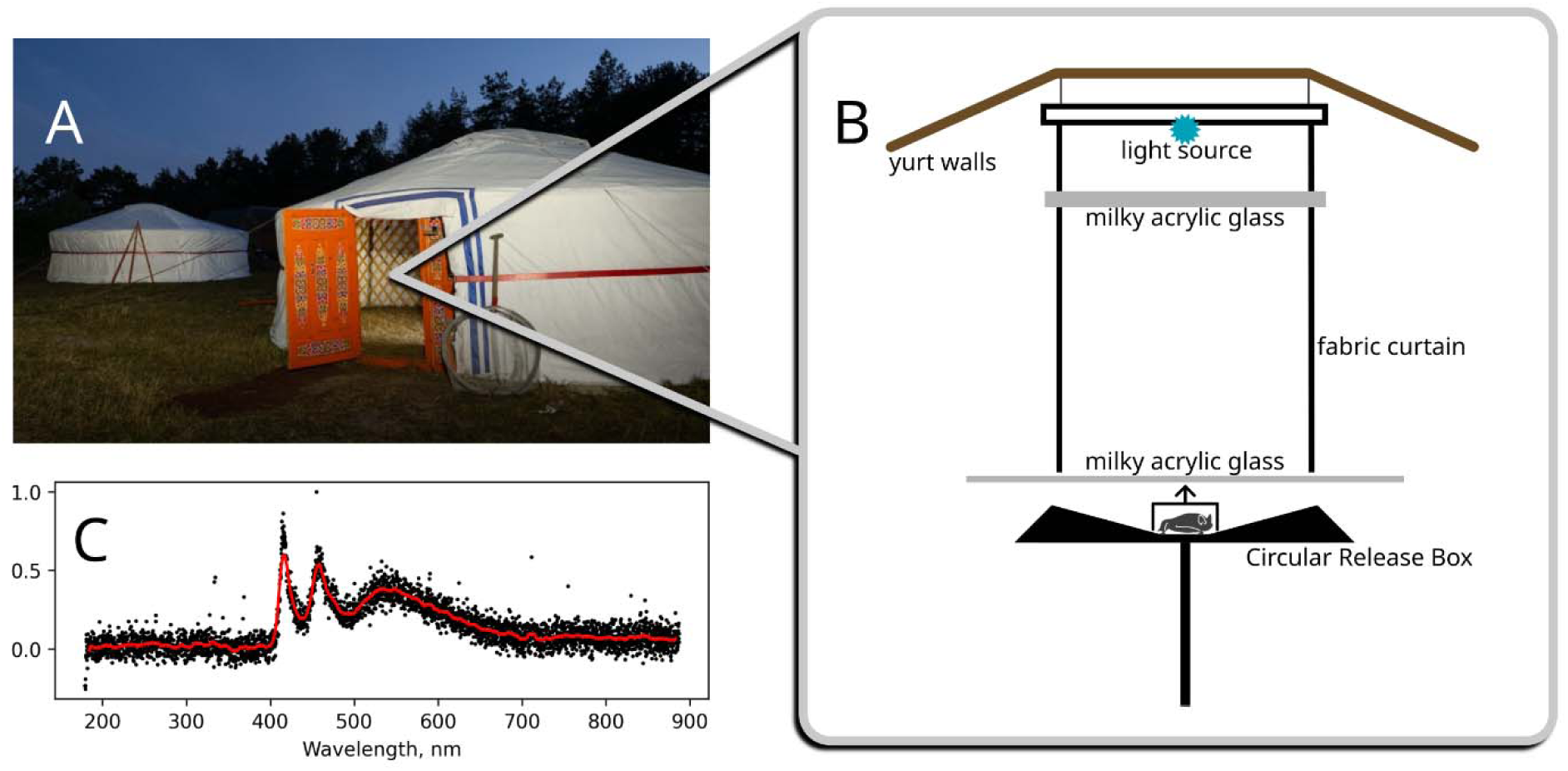
Study site, release light setup, and spectral characteristics of the light source. A) The Mongolian yurt served as a field laboratory for bat orientation tests. Prior to testing, bats were kept in acclimation boxes for the pre-exposure period, tuned for two conditions: light pre-exposure and darkness pre-exposure. B) Light setup inside the yurt was created to dissipate the light, centred above the circular release box, with two layers of milky acrylic glass, in order to avoid any directional cues for the tested animals. C) Normalized spectral radiance of the light source (in relative units). The same spectrum was used in both release and pre-exposure light setups.

Prior to testing, bats were placed in individual acclimation boxes and exposed to either starlight-level illumination or total darkness. The boxes were kept in a separate room in a chamber divided into two compartments: one held boxes with small windows of milky acrylic glass exposed to a light source, the other a larger carton box containing sealed, light-tight boxes.

Once inside the yurt, transfer of the bat from the acclimation box to the settling box took less than one minute and was carried out in darkness, the experimenter using night-vision goggles (PVS7 1×24 AlphaMod, S1550, Alpha Photonics GmbH & Co. KG, Germany) with a built-in infrared illuminator emitting at 940 nm (20 mW output). This wavelength lies well beyond the long-wavelength limit of chiropteran retinal sensitivity, which does not extend past approximately 680 nm [53], and bats have been reported not to respond behaviourally to far-red or infrared illumination [54]. Illumination at 940 nm therefore did not constitute a light exposure for the animals, including those acclimated to total darkness, and it falls outside the absorption bands of flavin in any redox form, so it cannot initiate cryptochrome photo-chemistry [2].

The bat was positioned in the middle of the arena and covered by a settling box (75 mm diameter, round 3D-printed in opaque black plastic, admitting no light). The side from which each bat was placed into the arena was randomised by selecting one of the four cardinal directions (N, S, W, E), to prevent any ob-server-side bias in the animals’ orientation.

After the settling, the bat was remotely released by a pulley system operated from outside the yurt, and was given 3 min to leave the arena by taking off. Take-off was detected acoustically from outside the yurt using a narrowband ultrasound detector (Pettersson D100), with the microphone mounted inside the yurt and headphones leading outside to the experimenter, following [17]; the onset of echolocation calls characteristic for free-flight indicated that the bat had become airborne. Take-off latency was recorded as the interval between release and the first detected call. Once a take-off had been detected, or once the 3 min had elapsed, the experimenter switched on conventional lighting inside the yurt before the evaluator entered, recaptured the bat, and recorded the departure heading from the tracks left by the bat on the arena surface. In two cases the track could not be read, and these individuals were excluded from the orientation analysis while being retained as departures in the comparison of latencies between groups. Between trials the chalk layer was removed and renewed, and the arena was rotated to a random orientation, so that any residual olfactory marks bore no fixed relationship to geographic direction.

Blinding was achieved by dividing experimental roles between two people. One selected the acclimation condition at random, transferred the bat to the yurt and prepared the arena; the other recorded take-off direction and was unaware of the condition under which the animal had been acclimated and tested.

### Light exposure and measurements

To ensure light provides no directional cue for the tested animals, we constructed the light dissipation setup inside a yurt. The light source was fixed at the top of the yurt, centred above the circular arena. In order to limit the illumination of the yurt walls, we put a fabric funnel to restrict light to the arena only. Before the animal could see the light, it was passing through two dissipation layers of milky acrylic glass: one immediately after the light source and another, the lid, before the arena itself (**Fig. 4B**).

To mimic the light conditions of a natural sky starry-night, the “starlight” condition, we used a combination of two LEDs: white (Avonec 3W star-plated LED CCT ca. 8000K) and UV (Avonec 3W star-plated LED 410-420 nm UV-A), to obtain a relatively flat spectrum across the relevant spectral band [55]. The spectral characteristics of the combined LEDs were measured with an Ocean Optics USB4000 fiber-optic spectrometer (Ocean Optics Inc., Dunedin, FL, USA) (**Fig. 4C**). Irradiance experienced by the animals was measured in the centre of the arena and inside the acclimation boxes with a Thorlabs PM400 optical power meter with a S130VC photodiode power sensor (Thorlabs, Inc., Newton, NJ, USA). We aimed to replicate the irradiance measured outdoors during a moonless night (E*_e_*= 31.6 µW/m^2^ ± 1.6 µW/m^2^, SD; corresponding to an illuminance of E*_v_* ≈ 2.8 mlx). We set up the light source to achieve an irradiance of E*_e_* = 35.5 µW/m^2^ ± 3.5 µW/m^2^ (SD; ca. 3.2 mlx) for acclimation boxes and E*_e_* = 24.1 µW/m^2^ ± 1.0 µW/m^2^ (SD; ca. 2.2 mlx) for the centre of the release box.

The “total darkness” condition inside the yurt was far below the sensitivity threshold of the optical power meter and verification was achieved in multiple steps. First, the condition inside the yurt was assessed by five independent human observers (2 during daytime, 3 at night). After dark adapting for 45 minutes inside the release room, none of the human observers was able to detect any light there. Second, night-time measurements were performed with sensitive light measurement devices inside the yurt at night during moonlit conditions. Here, a precision illuminance meter (ILT 5000 research radiometer with illuminance head SED-100, International Light Technologies, Inc., Peabody, MA, USA) and an all-sky camera system with a fisheye lens (Canon EOS6D, Canon Inc., Tokio, Japan; Sigma EXDG 8mm, Sigma Corporation, Kanagawa, Japan) was used to measure the luminance. The ILT5000 illuminance meter is NIST traceable with a measurement range of 0.15 mlx to 10^6^ lx with a cosine corrected detector head. The camera measures spatially resolved luminance [56,57] and has a detection limit of 0.005 mcd/m² and 0.01 mlx (ca. 0.1 µW/m²), respectively, when using the Sky Quality Software calibration (Euromix, Ljubljana, Slovenia). Inside the yurt at night, neither illuminance meter nor camera could detect any signal (**Fig. S2,** Electronic supplementary material 2). This is more than an order of magnitude below the visual detection threshold of *Myotis daubentonii* (0.3 mcd/m²) [58] which is on the order of humans [40,59]. Third, additional parallel measurements during daytime were performed using another precision illuminance meter (ILT1700, 0.8 mlx to 10^6^ lx range) outside of the yurt, as well as the aforementioned ILT5000 precision illuminance meter and the all-sky camera system inside of the yurt. These parallel daytime measurements (clear sky) allowed us to calculate how the yurt attenuates light like an optical filter. Cosine corrected horizontal illuminance was below the detection threshold of the ILT5000. Scalar Illuminance inside the yurt (but outside the arena) was measured to be E*_v,scal_* = 0.1 ± 0.01 mlx using the camera (**Fig. S3**, Electronic supplementary material 2), while outside a scalar illuminance of 100,000 lx was determined with the ILT1700 illuminance meter facing towards the sun. Inside the arena, the camera was not able to detect any signal (E*_v,scal_* < 0.01 mlx). This results in an attenuation factor of 10^-9^ inside the yurt but outside the arena and 10^-10^ inside the arena compared to the outside illuminance. During the experiment, the maximum nocturnal scalar illuminance was calculated to be 70 mlx [60] (89% moon illumination, 14° above horizon, negligible twilight and skyglow). Therefore, the resulting light level inside the yurt at night did not exceed 0.07 nlx (assuming 10^-9^ attenuation inside the yurt) or 7 plx respectively (assuming 10^-10^ attenuation inside the arena), which can be considered absolutely dark (“total darkness condition”). The acclimation boxes used for the total darkness pre-exposure were light-tight containers by construction, and no light could possibly penetrate. However, for completeness’ sake, we conducted daytime measurements analogous to the yurt attenuation. We used the ILT5000 illuminance meter inside and the ILT1700 illuminance meter outside of the box in parallel on a day with stable solar irradiance (clear sky). While outside again an illuminance of ca. 100,000 lx was determined, no signal was detected inside the box (i.e. below 0.15 mlx), which results in an attenuation of about 6.67 10^-8^. With the aforementioned maximum outdoor illuminance of 70 mlx, we calculate a maximum of 4.7 nlx scalar illuminance, establishing total darkness condifor the pre-exposure period as well.

### Statistical analysis

Data analysis and visualisation, with the exception of maximum-likelihood analysis, were performed in Python 3.12 [61]. Most circular-statistical tests were implemented using functions available in the Circular-Plotter repository (https://github.com/ftsellarius/Circular-Plotter). We used the Rayleigh test of uniformity to assess whether each sample showed significant orientation and the non-parametric Mardia–Watson–Wheeler (MWW) test for pairwise comparisons of samples [62]. Differences in angular concentration were tested using the test for equality of concentration parameters described by Mardia and Jupp [63].

The power of the Rayleigh test and the sensitivity of the MWW comparisons to a uniform alternative were evaluated in separate simulation analyses. For the Rayleigh-test power analysis, 10,000 samples of size n=25 were drawn from von Mises distributions with concentration parameters ranging from 0.25 to 2. Each sample was analysed using the Rayleigh test at α=0.05. For each concentration parameter, power was estimated as the proportion of simulated samples for which the null hypothesis of uniformity was rejected, and the type II error rate was calculated as 1-power. The procedure was repeated in 10 independent batches for each concentration value to assess Monte Carlo variability.

For the MWW sensitivity analysis, 10,000 samples were drawn from a uniform distribution and compared with the unchanged reference sample. We calculated the proportion of simulations yielding an MWW-test p-value equal to or greater than that observed in the corresponding comparison with the target sample. This value represents the probability, under the specified uniform alternative, of obtaining a result at least as non-significant as the observed result.

Within each experimental condition, the association between take-off direction and acclimation duration was tested using circular–linear correlation implemented in the Python package pycircstat2. This function evaluates the association between an angular and a linear variable and returns both a correlation coefficient and a p-value.

Take-off latency was compared among conditions using the Kruskal–Wallis test. Differences among conditions in the proportion of bats that failed to take off were tested using a Monte Carlo implementation of Fisher’s exact test on a 3x2 contingency table, followed by pairwise 2x2 Fisher’s exact test for control versus dark-acclimated and dark-tested sample. The Kruskal–Wallis and Fisher’s exact tests were implemented in SciPy [64].

Maximum-likelihood analysis was additionally performed using the CircMLE package in R [65].

## Supporting information

Electronic supplementary material 1

Electronic supplementary material 2

## Ethics

All work was conducted under the permit Nr.117.2024 to the Institute of Biology, University of Latvia. All experimental procedures were conducted in accordance with national and local guidelines for the use of animals in research.

## Data accessibility

The raw data and all code used for the analyses are available upon reasonable request.

## Authors’ contributions

FC: conceptualization, formal analysis, methodology, software, validation, visualization, writing—original draft, writing—review and editing; J.B.: investigation; writing—review and editing; K.S.: investigation, data curation, writing—review and editing; A.J.: conceptualization, methodology, data curation, investigation, resources, writing—review and editing, supervision; O.L.: conceptualization, data curation, investigation, project administration, resources, funding acquisition, writing—review and editing, supervision.

## Acknowledgments

We thank Dinora Abdulazhanova for supporting the equipment preparation and Olivia Leißring for her assistance with the experiments. We thank Oskars Keišs and Māris Jaunzemis for help with field installations. For help with bat catching, animal husbandry and yurt maintenance, we a grateful to Maria Janzen and Jake Elliott.

**Supplementary material 1**

CircMLE estimation results and Rayleight test power estimation

**Supplementary material 2**

Raw illuminance from all-sky camera system

## References

1. Lam BYK, Malkemper EP. 2026 Magnetosensation: the unsolved mystery. Physiological Reviews 106, 2383–2439. (doi:10.1152/physrev.00032.2025)

2. Hore PJ, Mouritsen H. 2016 The Radical-Pair Mechanism of Magnetoreception. Annual Review of Biophysics 45, 299–344. (doi:10.1146/annurev-biophys-032116-094545)

3. Xu J et al. 2021 Magnetic sensitivity of cryptochrome 4 from a migratory songbird. Nature 594, 535–540. (doi:10.1038/s41586-021-03618-9)

4. Blakemore R. 1975 Magnetotactic Bacteria. Science 190, 377–379. (doi:10.1126/science.170679)

5. Lefèvre Christopher T., Bazylinski Dennis A. 2013 Ecology, Diversity, and Evolution of Magnetotactic Bacteria. Microbiology and Molecular Biology Reviews 77, 497–526. (doi:10.1128/mmbr.00021-13)

6. Kirschvink JL, Gould JL. 1981 Biogenic magnetite as a basis for magnetic field detection in animals. Biosystems 13, 181–201. (doi:10.1016/0303-2647(81)90060-5)

7. Shaw J, Boyd A, House M, Woodward R, Mathes F, Cowin G, Saunders M, Baer B. 2015 Magnetic particle-mediated magnetoreception. Journal of The Royal Society Interface 12, 20150499. (doi:10.1098/rsif.2015.0499)

8. Holland RA, Kirschvink JL, Doak TG, Wikelski M. 2008 Bats use magnetite to detect the earth’s magnetic field. PLoS ONE 3, 1–6. (doi:10.1371/journal.pone.0001676)

9. Nimpf S et al. 2019 A Putative Mechanism for Magnetoreception by Electromagnetic Induction in the Pigeon Inner Ear. Current Biology 29, 4052–4059.e4. (doi:10.1016/j.cub.2019.09.048)

10. Nordmann GC, Balay SD, Kapuruge TN, Numi M, Leeb C, Nimpf S, Malkemper EP, Land-ler L, Keays DA. 2026 A global screen for magnetically induced neuronal activity in the pigeon brain. Science 391, 1155–1160. (doi:10.1126/science.aea6425)

11. Alcalde JT, Jimenez M, Brila I, Vintulis V, Voigt CC, Petersons G. 2020 Trans-continental 2,200 km migration of a Nathusius pipistrelle (Pipistrellus nathusii). Mammalia, 35–37.

12. Vasenkov D, Desmet J-F, Popov I, Sidorchuk N. 2022 Bats can migrate farther than it was previously known: a new longest migration record by Nathusius’ pipistrelle Pipistrellus nathusii (Chiroptera: Vespertilionidae). Mammalia, 1–3. (doi:10.1515/mammalia-2021-0139)

13. Lindecke O, Voigt CC, Pētersons G, Holland RA. 2015 Polarized skylight does not calibrate the compass system of a migratory bat. Biology Letters 11, 20150525. (doi:10.1098/rsbl.2015.0525)

14. Lindecke O et al. 2026 Disruptive effects of brief radiofrequency noise exposure on migratory bat navigation. Science 392, 977–979. (doi:10.1126/science.adq4418)

15. Holland RA, Thorup K, Vonhof MJ, Cochran WW, Wikelski M. 2006 Bat orientation using Earth’s magnetic field. Nature 444, 702–702. (doi:10.1038/444702a)

16. Wang Y, Pan Y, Parsons S, Walker M, Zhang S. 2007 Bats respond to polarity of a magnetic field. Proceedings of the Royal Society B: Biological Sciences 274, 2901–2905. (doi:10.1098/rspb.2007.0904)

17. Schneider WT, Holland RA, Keišs O, Lindecke O. 2023 Migratory bats are sensitive to magnetic inclination changes during the compass calibration period. Biol. Lett. 19, 20230181. (doi:10.1098/rsbl.2023.0181)

18. Netušil R, Tomanová K, Chodáková L, Chvalová D, Doležel D, Ritz T, Vácha M. 2021 Cryptochrome-dependent magnetoreception in a heteropteran insect continues even after 24 h in darkness. Journal of Experimental Biology 224, jeb243000. (doi:10.1242/jeb.243000)

19. Player TC, Hore PJ. 2019 Viability of superoxide-containing radical pairs as magnetoreceptors. The Journal of Chemical Physics 151, 225101. (doi:10.1063/1.5129608)

20. Zhang L, Malkemper EP. 2023 Cryptochromes in mammals: a magnetoreception misconception? Frontiers in Physiology Volume 14-2023.

21. Foley LE, Gegear RJ, Reppert SM. 2011 Human cryptochrome exhibits light-dependent magnetosensitivity. Nature Communications 2, 356. (doi:10.1038/ncomms1364)

22. Bassetto M, Reichl T, Kobylkov D, Kattnig DR, Winklhofer M, Hore PJ, Mouritsen H. 2023 No evidence for magnetic field effects on the behaviour of Drosophila. Nature 620, 595–599. (doi:10.1038/s41586-023-06397-7)

23. Engels S et al. 2014 Anthropogenic electromagnetic noise disrupts magnetic compass orientation in a migratory bird. Nature 509, 353–356. (doi:10.1038/nature13290)

24. Schwarze S, Schneider N-L, Reichl T, Dreyer D, Lefeldt N, Engels S, Baker N, Hore PJ, Mouritsen H. 2016 Weak Broadband Electromagnetic Fields are More Disruptive to Magnetic Compass Orientation in a Night-Migratory Songbird (Erithacus rubecula) than Strong Narrow-Band Fields. Frontiers in Behavioral Neuroscience Volume 10-2016.

25. Leberecht B et al. 2023 Upper bound for broadband radiofrequency field disruption of magnetic compass orientation in night-migratory songbirds. Proceedings of the National Academy of Sciences 120, e2301153120. (doi:10.1073/pnas.2301153120)

26. Munro U, Munro JA, Phillips JB, Wiltschko W. 1997 Effect of Wavelength of Light and Pulse Magnetisation on Different Magnetoreception Systems in a Migratory Bird. Australian Journal of Zoology 45, 189–198. (doi:10.1071/ZO96066)

27. Wiltschko W, Wiltschko R. 2007 Magnetoreception in birds: two receptors for two different tasks. Journal of Ornithology 148, 61–76. (doi:10.1007/s10336-007-0233-2)

28. Phillips JB, Diego-Rasilla FJ. 2022 The amphibian magnetic sense(s). Journal of Comparative Physiology A 208, 723–742. (doi:10.1007/s00359-022-01584-9)

29. Goforth KM, Lohmann CMF, Gavin A, Henning R, Harvey A, Hinton TL, Lim DS, Lohmann KJ. 2025 Learned magnetic map cues and two mechanisms of magnetoreception in turtles. Nature 638, 1015–1022. (doi:10.1038/s41586-024-08554-y)

30. Nordmann GC, Hochstoeger T, Keays DA. 2017 Magnetoreception—A sense without a receptor. PLOS Biology 15, e2003234. (doi:10.1371/journal.pbio.2003234)

31. Hore PJ. 2026 A hybrid compass mechanism combining radical pairs and magnetite crystals. Proceedings of the National Academy of Sciences 123, e2524093123. (doi:10.1073/pnas.2524093123)

32. Lindecke O, Holland RA, Pētersons G, Voigt CC. 2021 Corneal sensitivity is required for orientation in free-flying migratory bats. Communications Biology 4, 522. (doi:10.1038/s42003-021-02053-w)

33. Pētersons G. 2004 Seasonal migrations of north-eastern populations of Nathusius’ bat Pipistrellus nathusii (Chiroptera). Myotis 41–42, 29–56.

34. Lindecke O, Elksne A, Holland RA, Pētersons G, Voigt CC. 2019 Orientation and flight behaviour identify the Soprano pipistrelle as a migratory bat species at the Baltic Sea coast. Journal of Zoology 308, 56–65. (doi:10.1111/jzo.12654)

35. Muheim R, Bäckman J, Åkesson S. 2002 Magnetic compass orientation in European robins is dependent on both wavelength and intensity of light. Journal of Experimental Biology 205, 3845–3856. (doi:10.1242/jeb.205.24.3845)

36. Voigt CC, Roeleke M, Marggraf L, Pētersons G, Voigt-Heucke SL. 2017 Migratory bats respond to artificial green light with positive phototaxis. PLOS ONE 12, e0177748. (doi:10.1371/journal.pone.0177748)

37. Voigt CC, Rehnig K, Lindecke O, Pētersons G. 2018 Migratory bats are attracted by red light but not by warm-white light: Implications for the protection of nocturnal migrants. Ecology and Evolution 8, 9353–9361. (doi:10.1002/ece3.4400)

38. Lindecke O, Valerts D, Vintulis V, Schneider WT. 2025 Timing of circadian shifts in light/dark preference by migratory bats supports targeting of sunset for magnetic compass calibration. bioRxiv, 2025.10.02.680110. (doi:10.1101/2025.10.02.680110)

39. Spitschan M, Aguirre GK, Brainard DH, Sweeney AM. 2016 Variation of outdoor illumination as a function of solar elevation and light pollution. Scientific Reports 6, 26756. (doi:10.1038/srep26756)

40. Hecht S, Shlaer S, Pirenne MH. 1942 ENERGY, QUANTA, AND VISION. Journal of General Physiology 25, 819–840. (doi:10.1085/jgp.25.6.819)

41. Erkert HG. 2004 Extremely low threshold for photic entrainment of circadian activity rhythms in molossid bats (Molossus molossus; Chiroptera - Molossidae). Mammalian Biology 69, 361–374. (doi:10.1078/1616-5047-00158)

42. Stapput K, Thalau P, Wiltschko R, Wiltschko W. 2008 Orientation of Birds in Total Darkness. Current Biology 18, 602–606. (doi:10.1016/j.cub.2008.03.046)

43. Marhold S, Wiltschko W, Burda H. 1997 A magnetic polarity compass for direction finding in a subterranean mammal. Naturwissenschaften 84, 421–423.

44. Lohmann KJ. 1991 Magnetic Orientation By Hatchling Loggerhead Sea Turtles (Caretta Caretta). Journal of Experimental Biology 155, 37–49. (doi:10.1242/jeb.155.1.37)

45. Light P, Salmon M, Lohmann KJ. 1993 Geomagnetic Orientation of Loggerhead Sea Turtles: Evidence for an Inclination Compass. Journal of Experimental Biology 182, 1–10. (doi:10.1242/jeb.182.1.1)

46. Quinn TP. 1980 Evidence for celestial and magnetic compass orientation in lake migrating sockeye salmon fry. Journal of comparative physiology 137, 243–248. (doi:10.1007/BF00657119)

47. Myklatun A, Lauri A, Eder SHK, Cappetta M, Shcherbakov D, Wurst W, Winklhofer M, Westmeyer GG. 2018 Zebrafish and medaka offer insights into the neurobehavioral correlates of vertebrate magnetoreception. Nature Communications 9, 802. (doi:10.1038/s41467-018-03090-6)

48. Hiscock HG, Hiscock TW, Kattnig DR, Scrivener T, Lewis AM, Manolopoulos DE, Hore PJ. 2019 Navigating at night: fundamental limits on the sensitivity of radical pair magnetoreception under dim light. Quarterly Reviews of Biophysics 52, e9. (doi:10.1017/S0033583519000076)

49. Kattnig DR. 2026 Signals too small to sense: Physical and information-theoretic limits to in-duction-based magnetoreception in birds. (10.48550/arXiv.2602.23485)

50. Horton TW, Holdaway RN, Zerbini AN, Hauser N, Garrigue C, Andriolo A, Clapham PJ. 2011 Straight as an arrow: humpback whales swim constant course tracks during long-distance migration. Biology Letters 7, 674–679. (doi:10.1098/rsbl.2011.0279)

51. Beltran RS et al. In press. Lightscapes of fear: How mesopredators balance starvation and predation in the open ocean. Science Advances 7, eabd9818. (doi:10.1126/sciadv.abd9818)

52. Beltran RS, Yuen AL, Condit R, Robinson PW, Czapanskiy MF, Crocker DE, Costa DP. 2022 Elephant seals time their long-distance migrations using a map sense. Current Biology 32, R156–R157. (doi:10.1016/j.cub.2022.01.031)

53. Hope GM, Bhatnagar KP. 1979 Electrical response of bat retina to spectral stimulation: comparison of four microchiropteran species. Experientia 35, 1189–1191. (doi:10.1007/BF01963279)

54. Mistry S, McCracken GF. 1990 Behavioural response of the Mexican free-tailed bat, Ta-darida brasiliensis mexicana, to visible and infra-red light. Animal Behaviour 39, 598–599. (doi:10.1016/S0003-3472(05)80428-2)

55. Kolláth Z, Jechow A. 2023 Natural variation of the colour and spectrum of the night sky observed at a potential european reference site for dark skies. Journal of Quantitative Spectroscopy and Radiative Transfer 303, 108592. (doi:10.1016/j.jqsrt.2023.108592)

56. Jechow A, Kyba CCM, Hölker F. 2019 Beyond All-Sky: Assessing Ecological Light Pollution Using Multi-Spectral Full-Sphere Fisheye Lens Imaging. Journal of Imaging 5, 46. (doi:10.3390/jimaging5040046)

57. Kolláth Z, Cool A, Jechow A, Kolláth K, Száz D, Tong KP. 2020 Introducing the dark sky unit for multi-spectral measurement of the night sky quality with commercial digital cameras. Journal of Quantitative Spectroscopy and Radiative Transfer 253, 107162. (doi:10.1016/j.jqsrt.2020.107162)

58. Céchetto C, Jakobsen L, Warrant EJ. 2023 Visual detection threshold in the echolocating Daubenton’s bat (Myotis daubentonii). Journal of Experimental Biology 226, jeb244451. (doi:10.1242/jeb.244451)

59. Tinsley JN, Molodtsov MI, Prevedel R, Wartmann D, Espigulé-Pons J, Lauwers M, Vaziri A. 2016 Direct detection of a single photon by humans. Nature Communications 7, 12172. (doi:10.1038/ncomms12172)

60. Krisciunas K, Schaefer BE. 1991 A MODEL OF THE BRIGHTNESS OF MOONLIGHT. Publications of the Astronomical Society of the Pacific 103, 1033. (doi:10.1086/132921)

61. Van Rossum G, Drake FL. 2009 Python 3 Reference Manual. Scotts Valley, CA: Cre-ateSpace.

62. Batschelet E. 1981 Circular statistics in biology. London: Academic Press.

63. Mardia KV, Jupp PE. 2000 Directional statistics. Chichester; New York: J. Wiley.

64. Virtanen P et al. 2020 SciPy 1.0: Fundamental Algorithms for Scientific Computing in Python. Nature Methods 17, 261–272. (doi:10.1038/s41592-019-0686-2)

65. Fitak RR, Johnsen S. 2017 Bringing the analysis of animal orientation data full circle: model-based approaches with maximum likelihood. The Journal of Experimental Biology 220, 3878–3882. (doi:10.1242/jeb.167056)

