## Supplementary material for "Compass orientation of a migratory mammal in total darkness": Electronic supplementary material 1

We used the CircMLE [1] package in R to perform maximum-likelihood analysis of the samples and identify the most plausible underlying distribution. To estimate the false-negative probability of the Rayleigh test, we assessed its power using simulated samples drawn from a distribution with specified parameters. For samples with a concentration parameter matching that of the dark-acclimated group tested in darkness, the estimated power was 0.31, corresponding to a type II error rate of 0.69.

**Table S1. CircMLE estimation results.**The models with ΔAICc < 2 are in bold. Experimental conditions are denoted as acclimation and test condition, separated with a hyphen.

|  | **Control** | | **Light-darkness** | | **Darkness-darkness** | |
| --- | --- | --- | --- | --- | --- | --- |
| *Distribution name* | *ΔAICc* | *w* | *ΔAICc* | *w* | *ΔAICc* | *w* |
| Uniform | 5.348 | 0.02468 | 6.25 | 0.01813 | **0** | 0.38832 |
| Unimodal | **0** | 0.35777 | 2.572 | 0.11408 | **1.58** | 0.17627 |
| Symmetric modified unimodal | **0.065** | 0.34628 | **0** | 0.41283 | **1.679** | 0.16771 |
| Modified unimodal | 2.321 | 0.11208 | 2.619 | 0.11143 | 3.078 | 0.08332 |
| Homogenous symmetric bimodal | 8.224 | 0.00586 | 5.23 | 0.0302 | 3.915 | 0.05484 |
| Symmetric bimodal | 3.244 | 0.07065 | 2.454 | 0.12104 | 4.279 | 0.04571 |
| Homogenous axial bimodal | 3.764 | 0.05449 | 3.011 | 0.09163 | 4.724 | 0.0366 |
| Axial bimodal | 7.321 | 0.0092 | 5.054 | 0.03299 | 6.445 | 0.01547 |
| Homogenous bimodal | 6.121 | 0.01676 | 4.097 | 0.05324 | 6.112 | 0.01828 |
| Bimodal | 10.149 | 0.00224 | 6.705 | 0.01445 | 6.722 | 0.01348 |

**Figure S1. Power of the Rayleigh test across different concentration parameters for samples of size n = 25.** The vertical line indicates the concentration parameter estimated for the dark-acclimated group tested in total darkness.

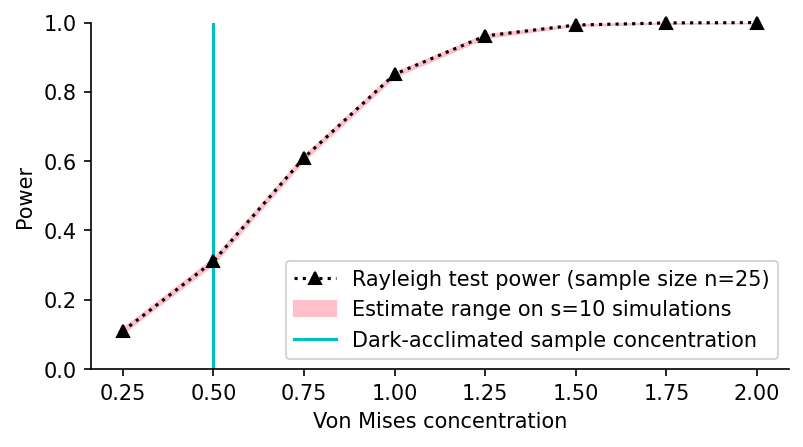

References

1.	Fitak RR, Johnsen S. 2017 Bringing the analysis of animal orientation data full circle: model-based approaches with maximum likelihood. *The Journal of Experimental Biology* **220**, 3878–3882. (doi:10.1242/jeb.167056)
