## Supplementary material for "Compass orientation of a migratory mammal in total darkness": Electronic supplementary material 2

All-sky camera system with a fisheye lens (Canon EOS6D, Canon Inc., Tokio, Japan; Sigma EXDG 8mm, Sigma Corporation, Kanagawa, Japan) was used to measure the illuminance inside the yurt.

**Figure S2. The results of the camera illumination measurements inside the yurt at night**

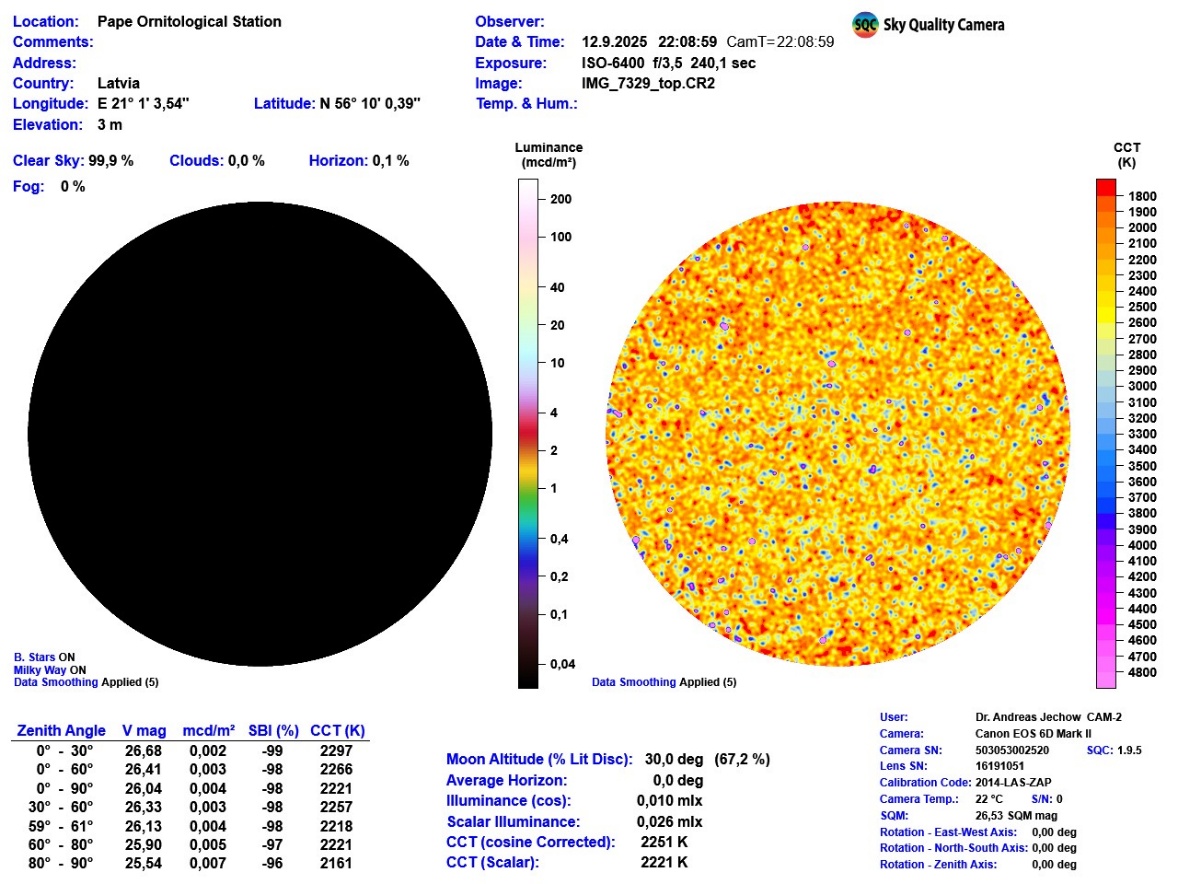

**Figure S3. The results of the camera illumination measurements inside the yurt at daytime.**

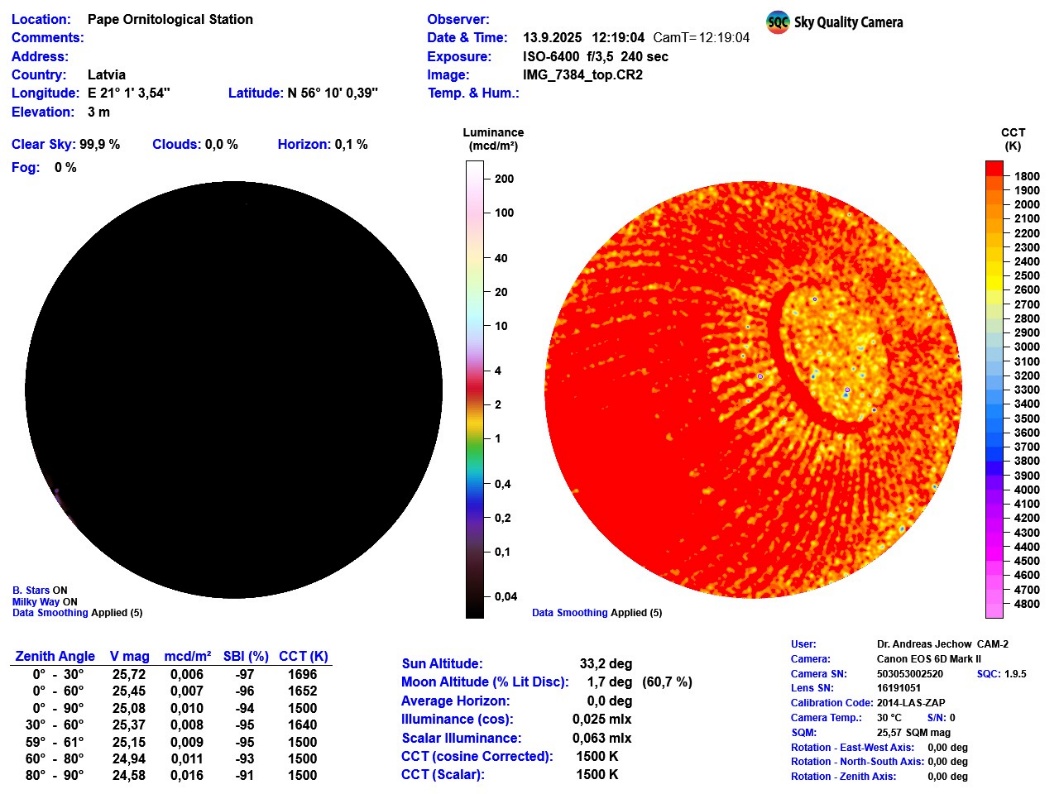
